# Differential chemoarchitecture of Purkinje neurons contributes to intrinsic firing properties

**DOI:** 10.1101/2021.01.28.428615

**Authors:** Cheryl Brandenburg, Lindsey A. Smith, Michaela B.C. Kilander, Morgan S. Bridi, Yu-Chih Lin, Shiyong Huang, Gene J. Blatt

**Author notes:** Corresponding author (CB), Address: Hussman Institute for Autism, 801 W Baltimore St. Suite 301, Baltimore, MD 21201. Senior authors (SH) (GJB).

## Abstract

Purkinje cells (PCs) are central to cerebellar information coding and appreciation for the diversity of their firing patterns and molecular profiles is growing. Heterogeneous subpopulations of PCs have been identified that display differences in intrinsic firing properties without clear mechanistic insight into what underlies the divergence in firing parameters. Although long used as a general PC marker, we report that the calcium binding protein parvalbumin labels a subpopulation of PCs with a conserved distribution pattern across the animals examined. We trained a convolutional neural network to recognize the parvalbumin-positive subtype and create maps of whole cerebellar distribution and find that PCs within these areas have differences in spontaneous firing that can be modified by altering calcium buffer content. These subtypes also show differential responses to potassium and calcium channel blockade, suggesting a mechanistic role for variability in PC intrinsic firing through differences in ion channel composition. It is proposed that ion channels drive the diversity in PC intrinsic firing phenotype and parvalbumin calcium buffering provides capacity for the highest firing rates observed. These findings open new avenues for detailed classification of PC subtypes.

## Introduction

The cerebellum’s repeated and precise geometric arrangement has provided vast insight into the complex relationships between neural circuits, plasticity and behavior. Its signature stereotyped circuitry was carefully dissected by Sir John Eccles and led to his seminal work “The Cerebellum as a Neuronal Machine” (Eccles, J.C. et al., 1967) and to his statement, “It seems likely that the cerebellum may be the first fragment of the higher levels of the nervous system to be understood in principle, all the way from peripheral input to peripheral output” (Eccles J.C., 1973). This optimistic prediction for a complete computational model of cerebellar influence on behavior has been challenged by the fact that, underlying this apparent uniform structure, is a complicated molecular code that segregates the cerebellum into an intricate topography of transverse zones, parasagittal stripes and microzones that are highly conserved between individuals and throughout evolution (for review see: Sotelo, 2020; Apps et al., 2018; Cerminara et al., 2015; Apps and Hawkes, 2009; Sillitoe and Joyner, 2007).

Purkinje cells (PCs) act as a template around which cerebellar circuit architecture is built (White and Sillitoe, 2013; Sotelo, 2004) and demarcate zones and stripes based on a number of molecular markers that correlate with afferent and efferent topography (Voogd et al., 2003; Sugihara and Shinoda 2004, 2007). Heterogeneous populations of PCs in parasagittally-oriented stripes were described by Gravel et al. (1987) and are perhaps best exemplified by labeling of aldolase C (originally zebrin II), which has served as a useful landmark to compare other markers that appear complementary (Brochu et al., 1990; Fujita et al., 2014). As aldolase C positive and negative stripes are differentially contacted topographically by cells projecting from the inferior olive and have been shown to have differences in expression of the glutamate transporter EAAT4 (Nagao et al., 1997; Dehnes et al., 1998; Hawkes, 2014), it stood to reason that these groups of PCs would have functional physiological differences. Only recently have differences in physiological properties between PC stripes been reported, notably that aldolase C negative PCs display higher simple spike firing frequencies than positive cells (Xiao et al., 2014; Zhou et al., 2014; Nguyen-Minh et al., 2018). However, the underlying mechanisms for these differences are unresolved because there is no evidence that aldolase C itself is responsible for the differences and researchers have pointed to the need for additional markers to help clarify the divergence in physiological parameters (Apps et al., 2018).

Here, we report that the calcium binding protein parvalbumin (PV) is not uniformly expressed throughout the cerebellum, but rather labels a subpopulation of PCs that cut across aldolase C stripes. In addition to calbindin, PV has been extensively employed as a general PC marker due to numerous reports of ubiquitous expression in the cerebellum (Baimbridge and Miller, 1982; Endo et al., 1985; Christakos et al.; 1987; Rogers, 1989; Scotti and Nitsch, 1992; Celio, 1990; Bastianelli, 2003; Schwaller et al., 2002, Whitney et al., 2008). However, one report in primate cerebellum observed a proportion of PV negative cells (Fortin et al., 1998) with a similar finding in avian cerebellum (Wylie et al., 2011). We show that, like markers for stripes, PV has a distinct pattern of expression that is conserved between individual animals, but is unlike the striping pattern observed for aldolase C.

This surprising distribution pattern of parvalbumin in the mouse cerebellum further divides lobules into heterogeneous PC populations and provides multiple novel avenues for exploration to understand the mechanisms responsible for differential firing between PCs and modules. PV is known to contribute to a cell’s capacity for high firing rates through its function as a calcium binding protein (e.g. Albéri et al., 2013; Rainnie et al., 2006). Additionally, basket neurons have been found to shift between low PV and high PV states, which is thought to be important in hippocampal plasticity (Donato et al., 2013). We hypothesized that the known association of PV with higher spike rates may explain the rate differences observed between PC subtypes and recorded spontaneous firing frequencies of PCs from areas of high intracellular PV expression and from areas that lack PV expression to compare populations. PV positive (+) PCs are more likely to display burst-pause patterns instead of tonic firing and to have a higher spontaneous firing rate than PV negative (-) cells, which often display tonic firing without bursting. We found that mimic of PV with a similar calcium buffer leads to increased firing rates and pausing time in PV-cells, PV knock-out mice have PCs with decreased firing rates and that blocking potassium and calcium channels differentially alter the burst-pause dynamics observed in PC subtypes.

We find that the diverse firing phenotypes displayed by PCs can be shifted into distinct patterns with ion channel block. Since burst-pause phenotypes are known to be controlled by ion channels (Llinas and Sugimori, 1980; Raman and Bean, 1997, 1999; Nam and Hockberger, 1997; Womack and Khodakhah, 2002; Shim et al., 2018), it seems practical that intrinsic firing properties such as bursting and firing rate would be modified by differential ion channel expression and/or function.

Taken together, regulation of intracellular calcium both by ion channel function and parvalbumin expression may be an important driver of the diverse spontaneous firing phenotypes seen across individual PCs. By adding to the repertoire of available markers that can segregate PCs into subtypes, finer mapping of cerebellar microcircuitry can be achieved. Improved understanding of cerebellar microcircuitry can lead to more detailed descriptions of physiological parameters responsible for appropriate information coding to the deep cerebellar nuclei and thereby enhance understanding of cerebellar function.

## Methods

### Animals

C57BL/6 mice were purchased from the Comparative Medicine/Veterinary Resources at the University of Maryland School of Medicine (Baltimore, MD, USA). PV-tdTomato mice (Stock #: 027395) and Pvalb-/- (Stock #: 027503) were purchased from the Jackson Laboratory and offspring were produced from heterozygous breeding pairs. Animals were housed in the animal facility with free access to food and water on a 12/12 h light/dark cycle. All experiments involving animal procedures were approved by the Institutional Care and Use Committees (IACUC) of the University of Maryland School of Medicine and the Hussman Institute for Autism.

### Immunohistochemistry

Following flush with PBS until the liver was pale, animals were transcardialy perfused with 80mL of 4% paraformaldehyde (PFA) over the course of twenty minutes. Brains were removed and placed in 4% PFA for 12 hours at 4°C before cryoprotection in 10% then 30% sucrose until sunk. Cryostat cut sections were stored in a cryoprotectant solution until use. In some cases, perfused brains were cut on a vibratome (Leica VT1000S) without cryoprotection.

PCs are surrounded by dense parvalbumin innervation from nearby basket cells, the optimal dynamic range of antibody concentration was critical to clearly observe PV negative cells, as described in Hoffman et al. (2016). For Ni^2+^ DAB, serial 40µm thick sections (every 6^th^) throughout the entire cerebellum of four week old C56/Bl6 mice (the pattern was also confirmed in one P21 mouse-data not shown) were rinsed of glycerol cryoprotectant three times for two minutes in a scientific microwave (Ted Pella) at 35 degrees and 150 watts (all following rinses were performed this way). Sections were blocked with 8% donkey serum in TBST for thirty minutes then incubated in 8% donkey serum TBST and primary antibody (1:4,000 guinea pig parvalbumin, GP72 Swant) for one hour at room temperature then 47 hours at 4°C followed by a rinse, one hour in secondary antibody (Jackson anti-guinea pig 1:600), another rinse, one hour in 1:500 A/B solution (Vector), rinse, twenty minutes in DAB solution (95mM nickel (II) sulfate hexahydrate (Sigma 106727); 0.55mM 3,3′-diaminobenzidine tetrahydrochloride hydrate (Sigma 32750) and 3% (v/v) hydrogen peroxide in TBS) and a final rinse before alcohol dehydration, xylene and coverslipping with DPX.

Fluorescence immunohistochemistry was performed in the same manner with increased PV primary concentration (guinea pig parvalbumin 1:2,000 GP72 Swant; mouse parvalbumin 1:2000 (or 1:300 where indicated) Sigma P3088; mouse calbindin 1:3,000 Sigma C9848; aldolase C 1:100 alexa fluor 647 Santa Cruz sc-271593), but after rinsing sections were incubated in secondary antibodies (donkey anti-guinea pig 555 1:1,000 Sigma SAB4600298; goat anti-mouse 546 1:400 Invitrogen A-11030; goat anti-mouse 488 Invitrogen 1:400 A-11001) for three hours, rinsed, dehydrated in alcohol, xylene and coverslipped with DPX.

### Fully automated distribution maps of parvalbumin positive cell types

#### A. Image Processing

Coronal Ni^2+^ DAB stained serial sections were imaged with a Microbrightfield (MBF) Zeiss, Stereoinvestigator system throughout the entire cerebellum in each of five mice. Image stacks were collapsed into deep focus files (MBF) at a resolution of 0.25 um/pixel. Slices were sectioned at 40µm and imaged in 20µm stacks at 1µm intervals. The digitized images were then uploaded to Aiforia™ Cloud image processing and management platform (Aiforia Technologies, Helsinki, Finland) for analysis with deep learning convolutional neural networks (CNN) and supervised learning.

#### B. AI Model Training Parameters

A supervised, multi-layered, CNN was trained on annotations from digitized images to recognize multiple DAB positive cell types using the cloud-based Aiforia™ platform. The algorithm was trained on the most diverse and representative images for this data set. 17 images (17/128= 13% of total dataset) constituted training data. When teaching AI, representativeness of the training slides is more critical than the number of training slides. Therefore, diverse training data were included to capture the variability in image/staining quality across the entire data set. We also chose slides with known artifacts and trained the AI model to exclude them from analysis as background.

The AI model consisted of multiple feature layers, containing unique classes that were annotated for CNN input data. The AI model consisted of four feature layers: 1) Tissue segmentation 2) Purkinje and molecular layer tissue sub region segmentations 3) DAB positive object detector for 3 subclasses within the Purkinje layer segmentation (parvalbumin positive Purkinje cells, parvalbumin negative Purkinje cells and interneurons) 4) DAB positive interneurons were identified using an object detector within the molecular layer segmentation. Individual CNNs were trained for each layer using the image augmentation parameters, perceptive view (field of view), and level complexity summarized in Table 1. All four layers were merged into a chained analysis pipeline, where segmentation results from the first layer are used as a cropping mask in the next layer, and so on, to detect and quantify the number of DAB positive cells within either Purkinje or Molecular layers across total tissue.

**Table 1.** CNN Image Augmentation Parameters, Field of View, and Complexity

|  | **Layer 1:**<br>**Tissue**<br>**segmentation** | **Layer 2:**<br>**Purkinje and**<br>**Molecular layer**<br>**segmentation** | **Layer 3:**<br>**Purkinje layer**<br>**cells** | **Layer 4:**<br>**Molecular**<br>**layer cells** |
| --- | --- | --- | --- | --- |
| **Scale (max/min)** | -10/10 | -10/10 | -10/10 | -1/1.01 |
| **Aspect Ratio** | 10 | 10 | 10 | 1 |
| **Maximum Shear** | 10 | 10 | 10 | 1 |
| **Luminance**<br>**(max/min)** | -10/10 | -10/10 | -10/10 | -1/1.01 |
| **Contrast (max/min)** | -10/10 | -10/10 | -10/10 | -1/1.01 |
| **Max. white balance**<br>**change** | 5 | 5 | 5 | 1 |
| **Noise** | 2 | 2 | 2 | 0 |
| **Field of View** | 50.4μm | 13.8 μm | | |
| **Complexity** | Intermediate | Extra Complex | Complex | Intermediate |

The ground truth, or features used to train the AI model, was annotated for each layer within the cloud-based Aiforia™ platform and constituted input data for each CNN. The first feature layer was annotated using semantic segmentation to distinguish the total tissue from the glass slide. The second feature layer was annotated using semantic segmentation to distinguish the Purkinje cell layer from the molecular layer within total tissue. The ground truth for the third layer utilized an object detector with an 18µ m diameter for parvalbumin positive Purkinje cells and parvalbumin negative Purkinje cells, with 10µ m for interneurons within the Purkinje layer. The ground truth for the fourth feature layer utilized an object detector with a 10µ m diameter to label interneurons within the Molecular Layer. Features that were considered artifact (glass slide, debris, out of focus regions) were annotated as background, and constituted additional input training data for the multi-layered CNN.

### Ex vivo slice electrophysiology

Cerebellar slices were prepared from P25-P45 C57/Bl6 mice and P30-P36 Pvalb-/- (modified from Eguchi et al., 2020). Mice were anesthetized with isoflurane and brains were removed in warm (34°C) dissecting artificial cerebrospinal fluid (aCSF) containing (in mM): 210 sucrose, 2.5 KCl, 1.25 NaH2PO4, 26 NaHCO3, 10 D-glucose, 0.5 ascorbic acid, 2 sodium pyruvate, 3 myo-inositol, 1 kynurenic acid, 4 MgCl2 and 0.1 CaCl2 saturated with 95% O2 and 5% CO2. Coronal sections were cut at 300µm on a vibratome (Leica VT1200s) in dissecting aCSF before being transferred to room temperature aCSF containing (in mM): 126 NaCl, 2.5 KCl, 1 NaH2PO4, 24 NaHCO3, 10 D-glucose, 2 MgCl2 and 2.5 CaCl2 saturated with 95% O2/5% CO2 and held for at least one hour before recording.

The loose-patch configuration in voltage clamp mode (zero current injection) was used to reliably obtain recordings of spontaneous firing by minimizing disturbance of the cell membrane (Perkins, 2006) with borosilicate glass pipettes that had an open pipette resistance of 2-4Ω with slice maintained near 32°C. The drugs NBQX (10µM), DL-AP5 (50 µM) and picrotoxin (100 µM) were added to pharmacologically block any synaptic contribution to PC firing. Recordings were collected using an Axon MultiClamp 700B amplifier (Molecular Devices) filtered at 10 kHz and digitized at 25 kHz with a National Instruments 150 digital-to-analog converter under the control of Igor Pro software (WaveMetrics, v6.37, 151). Spontaneous (intrinsically generated) firing rates were quantified over five minutes. Addition of 150µM EGTA/AM (Atluri & Regehr, 1996), 1mM TEA (McKay & Turner, 2004) or 500 nM ω-Conotoxin MVIIC (McDonough et al., 1996) was achieved through bath application. TEA had effects in less than fifteen minutes, while a minimum of forty minutes incubation was necessary before recordings of EGTA and ω-Conotoxin MVIIC began, as reported in the referenced studies. Analysis of spike rate over time was achieved through generation of histograms with the Igor software add-on, NeuroMatic (Rothman & Silver, 2018). After tests for normality, either a Student’s t-test or a Mann-Whitney test was utilized to compare differences in mean intrinsic properties between groups, while a paired t-test or Wilcoxon matched-pairs signed rank test was used to compare paired cells before and after drug application with Graph Pad Prism version 8.0.

### Immunoblotting

Cerebellar slices were frozen at −80°C immediately after electrophysiological recordings. Slices from each individual animal were combined in separate pools and homogenized on ice in RIPA buffer (Cell Signaling Technology) containing 1% Halt™ protease and phosphatase inhibitor cocktail (Thermo Scientific) and 1mM PMSF with a handheld Pellet Pestle® cordless motor (Kimble Chase). Homogenates were briefly sonicated, rotated for 1 hour at 4°C and centrifuged at 14000 x g in a cold microfuge. Supernatant was collected and total protein concentration was determined using BCA assay (Thermo Scientific). 20ug of each homogenate was mixed with 2 x Laemmli sample buffer (BioRad) containing 100mM DTT and denatured at 95°C for 5 min. Proteins were separated according to size by SDS-PAGE in a Mini-Protean TGX stain-free 4-20% gradient gel (BioRad) and thereafter transferred onto a PVDF membrane. The blot was blocked in 5% non-fat milk in TBS-T and probed with primary antibodies mouse monoclonal anti-parvalbumin (Sigma-Aldrich Cat# P3088, 1:2000) and mouse monoclonal anti-β-actin (Sigma-Aldrich Cat# A5441, 1:10000) o/n at 4°C. Subsequently, blots were incubated with secondary HRP-conjugated goat anti-mouse IgG antibodies (Thermo Fisher Scientific Cat# G-21040, 1:10000). Chemiluminescence signal was generated using Clarity ECL substrate (BioRad) and detected with a ChemiDoc (BioRad) at Optimal Auto-Exposure setting. Images of SDS-PAGE gel, as well as unstained PVDF membrane, were acquired pre and post transfer procedures to evaluate uniformity of sample loading and efficiency of electrophoretic transfer of proteins. PageRuler™ Plus Prestained Protein Ladder (Termo Scientific) was used as a reference for protein size determination.

## Results

### Differential chemoarchitecture of Purkinje neurons

Although PV is commonly used as a general PC marker, its expression is limited to distinct sub regions within cerebellar lobules (Figure 1). Interneurons in the molecular layer show intense PV staining equally throughout each region, but the PC dendritic trees that comprise the molecular layer show alternating intensity of PV staining. Upon closer examination, dendritic areas with lower PV intensity are situated above PCs that do not contain intracellular PV. However, they are surrounded by a “ring” of PV, which is innervation from interneurons that contact PC soma and form the pinceau (Ramon y Cajal, 1911, 1995 (translated); Palay and Palay, 1974). When high concentrations of primary antibody are used, this ring can become thicker and may be mistaken for intracellular stain, which may be a reason these patterns have been overlooked in previous reports. Therefore, optimal dynamic range of antibody concentrations (Hoffman et al., 2016) are important for identification of PC subtypes. There are examples in the literature of images where PV is not staining intracellularly, however, it is not discussed as the studies were not focused on PV (Jeong et al., 2000; Lee et al., 2018, Zhao et al., 2011). PCs not labeled with PV do stain for calbindin, confirming that the PV-PCs are labeled intracellularly by other markers (Figure 2).

**Figure 1.**
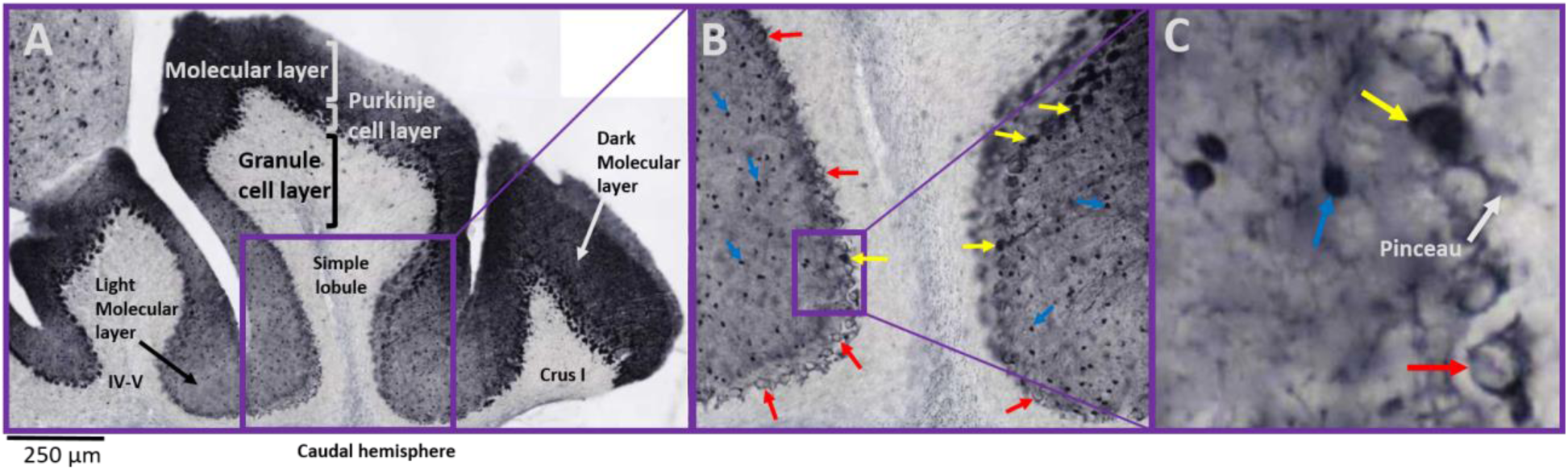
Parvalbumin labels a subpopulation of Purkinje cells. A) A coronal section stained with Ni^2+^ DAB parvalbumin guinea pig (Swant) shows alternating areas of parvalbumin-positive and -negative Purkinje cells. The dendritic trees of Purkinje cells in the molecular layer are also more darkly stained above the positive cells. B) The first inset shows a magnified view where red arrows indicate parvalbumin-negative and yellow arrows indicate parvalbumin-positive Purkinje cells. Interneuron arrows are blue. C) The second inset magnifies further to show individual cell types. Notice that interneurons are intensely parvalbumin-positive, which creates a “ring” of label around parvalbumin-negative Purkinje cells from basket cell somatic contact and pinceau formation, while positive Purkinje cells stain darkly throughout the cytoplasm.

**Figure 2.**
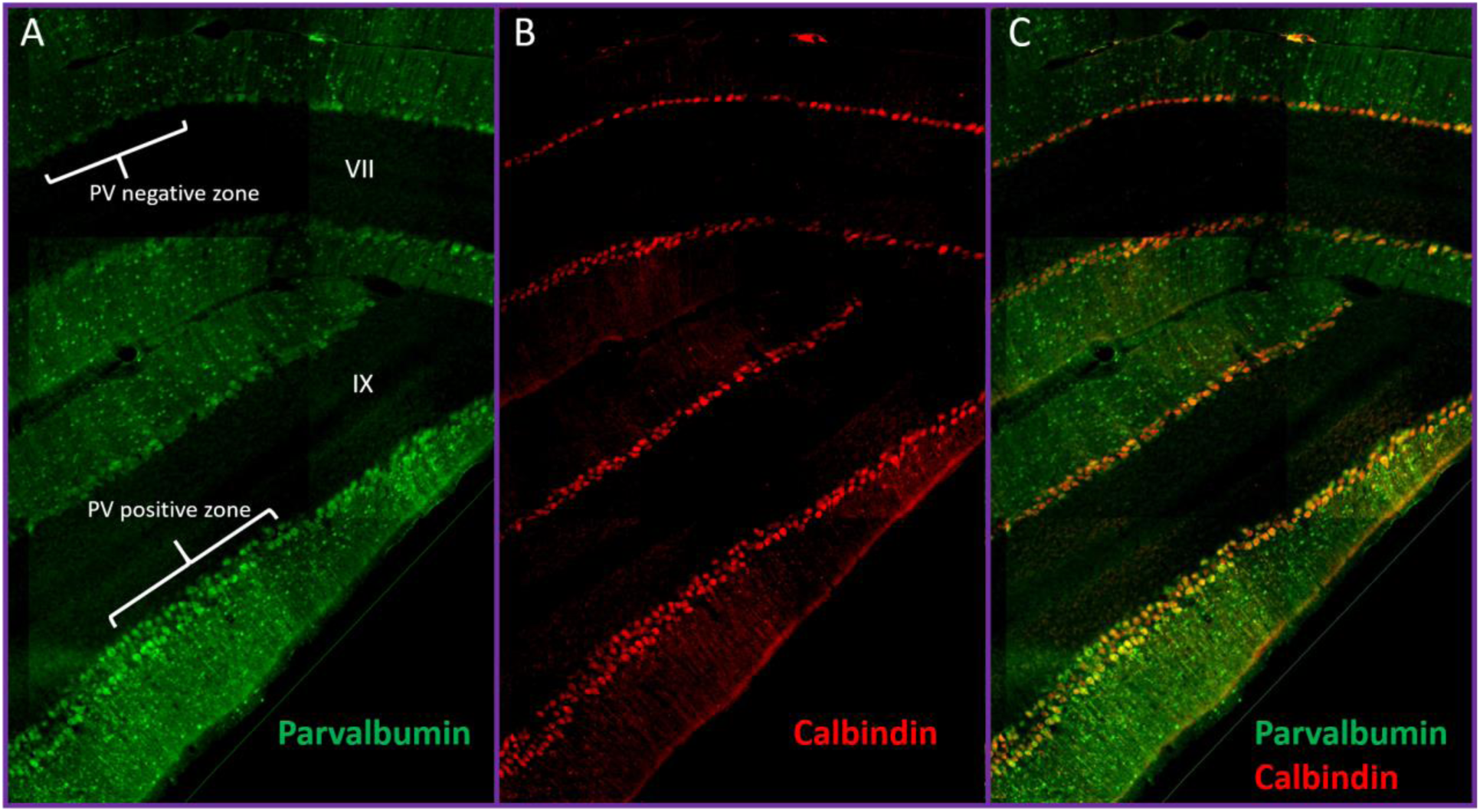
Parvalbumin-negative zones of Purkinje cells contain calbindin. A) Sections adjacent to those that had been stained with Ni2+ DAB showed the same pattern of parvalbumin-positive zones segregating to the perimeter of the cerebellum (PV+ zone in brackets) with negative zones more interior (PV-zone in brackets) only having surrounding interneuron input. B) To confirm that PV-zones stain with other Purkinje cell markers, calbindin was labeled and appears homogeneous throughout each region. C) The PV+ zone shows yellow cells, indicating colocalization of PV (guinea pig Swant) and calbindin, while the PV-zone is only staining for calbindin (red) intracellularly. Areas of the vermis are indicated.

An alternative explanation for the segregation of PV+ PCs to the perimeter of the cerebellum is poor perfusion in the interior of the sections. However, intensity of PV in interneurons is equal throughout the sections, PV+ PC white matter tracts are evident throughout, the deep cerebellar nuclei are appropriately stained with PV and PCs that are PV-stain with calbindin. Fluorescent Nissl positivity does not show any gradients across sections and reveals cell bodies of PCs that have equal intensity in both PV+ and PV-areas (Figure 3).

**Figure 3.**
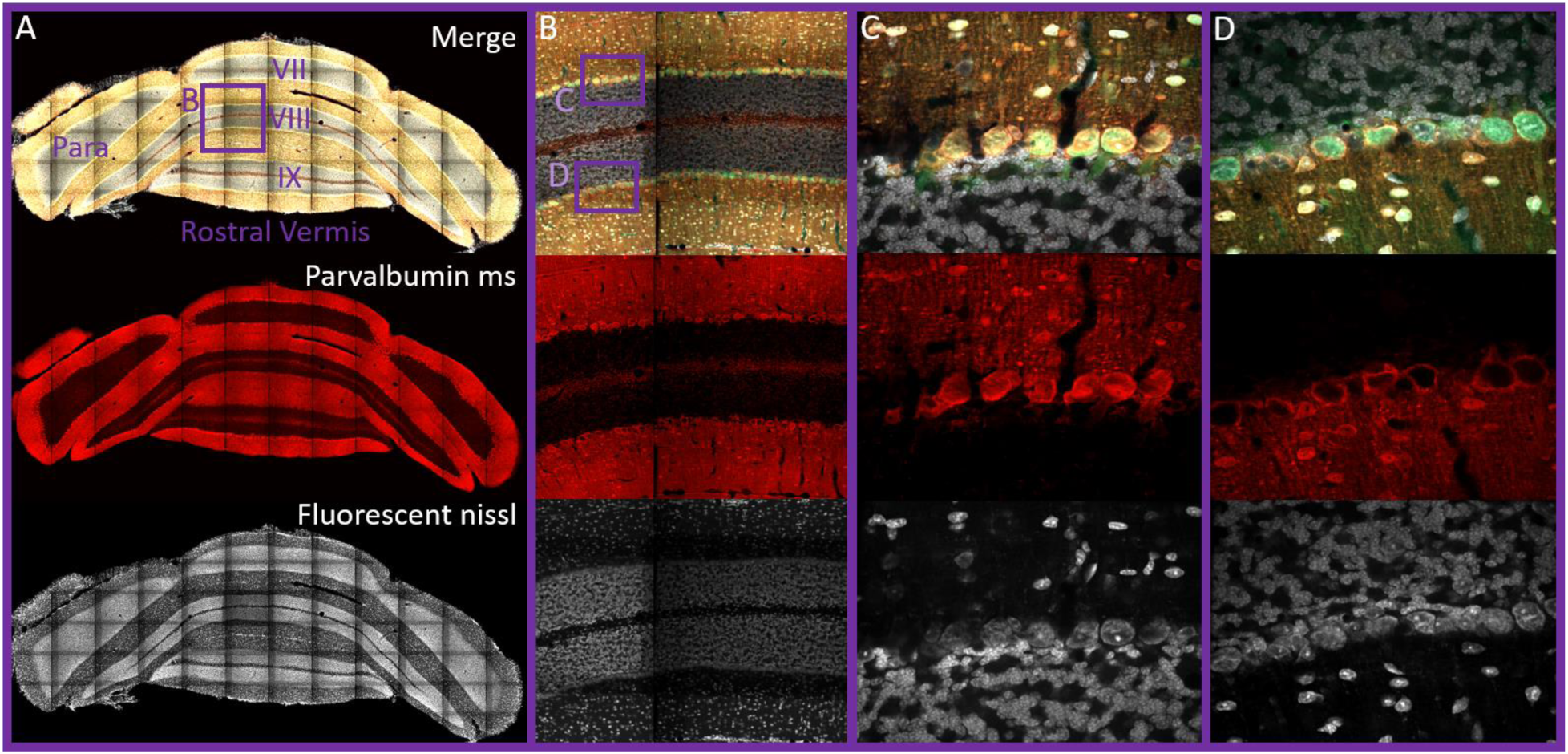
Dilution of parvalbumin antibody can affect intracellular staining appearance. A) Macroscopic view of the rostral cerebellum reveals even staining across lobules with fluorescent Nissl, displaying proper perfusion and all Purkinje cell bodies. Intracellular parvalbumin expression with fluorescence is difficult to appreciate from a macroscopic view, but shows that general labeling for interneurons and white matter tracts is even throughout the section. B) Inset from A showing two rows of Purkinje cells in the vermis that display mostly parvalbumin-positive Purkinje cells (upper) (C) and mostly parvalbumin-negative Purkinje cells (lower) (D). Panel A and B are taken at 20x magnification and panels C and D are taken at 63x magnification on a Zeiss confocal spinning disc. Parvalbumin mouse (ms) (Sigma) staining replicates the same pattern shown with the parvalbumin guniea pig antibody (Swant Figure 2) and all Purkinje cells have equal Nissl staining, suggesting differential antibody labeling from different companies, angle of cut of the cell and perfusion artifacts are not likely explanations for differences. Areas of the Vermis are labeled as well as the Paramedian lobule (Para).

Since PV did not appear to follow the typical striping pattern, a new distribution map was necessary to understand the overall topography of PV+ PCs across lobules. Furthermore, it was unclear whether these patterns would be conserved across individual animals (as aldolase C stripes are) or whether populations would shift since PV expression has been reported to be activity-dependent in other brain regions (Philpot et al., 1997; Patz et al., 2004; Kinney et al., 2006; Chaudhury et al., 2008; Mix et al., 2014). Therefore, serial sections throughout the entirety of the cerebellum of five mice were stained for PV (as in Figure 1 and supplemental figure 1) and a subset of these sections were used to train a supervised, multi-layered, convolutional neural network (Aiforia™). This AI model was able to accurately identify PV+ cells and PV-cells (based on the ring of input around the soma) to produce images overlaid with color coded cell types (Figure 4A). When adding all PCs counted in each animal across serial sections and dividing by the total area of the Purkinje layer, PV+ PCs represented 60.3% of the total population, with PV-cells as the remaining 39.7% (Figure 4B). As these are whole cerebellum counts, percentages may vary when comparing vermis to cerebellar hemispheres or individual lobules. Serial sections analyzed throughout one full cerebellum can be found in supplemental figure 1, which includes PV+ interneurons. These distribution maps show that PV+ PCs segregate to the perimeter of each section (corresponding to the surface of the cerebellum) across each lobule in the hemisphere as well as the vermis. Specifically, the yellow circles that represent PV+ PCs are the most lateral portion of each lobule and the most superior or inferior portions of the vermis, while the medial PCs of each region are mainly PV-PCs (red circles). The PV topography is therefore another conserved distribution pattern, as the patterns are the same across each animal examined (n=5). The pattern is best appreciated by examining supplemental figure 1.

**Figure 4.**
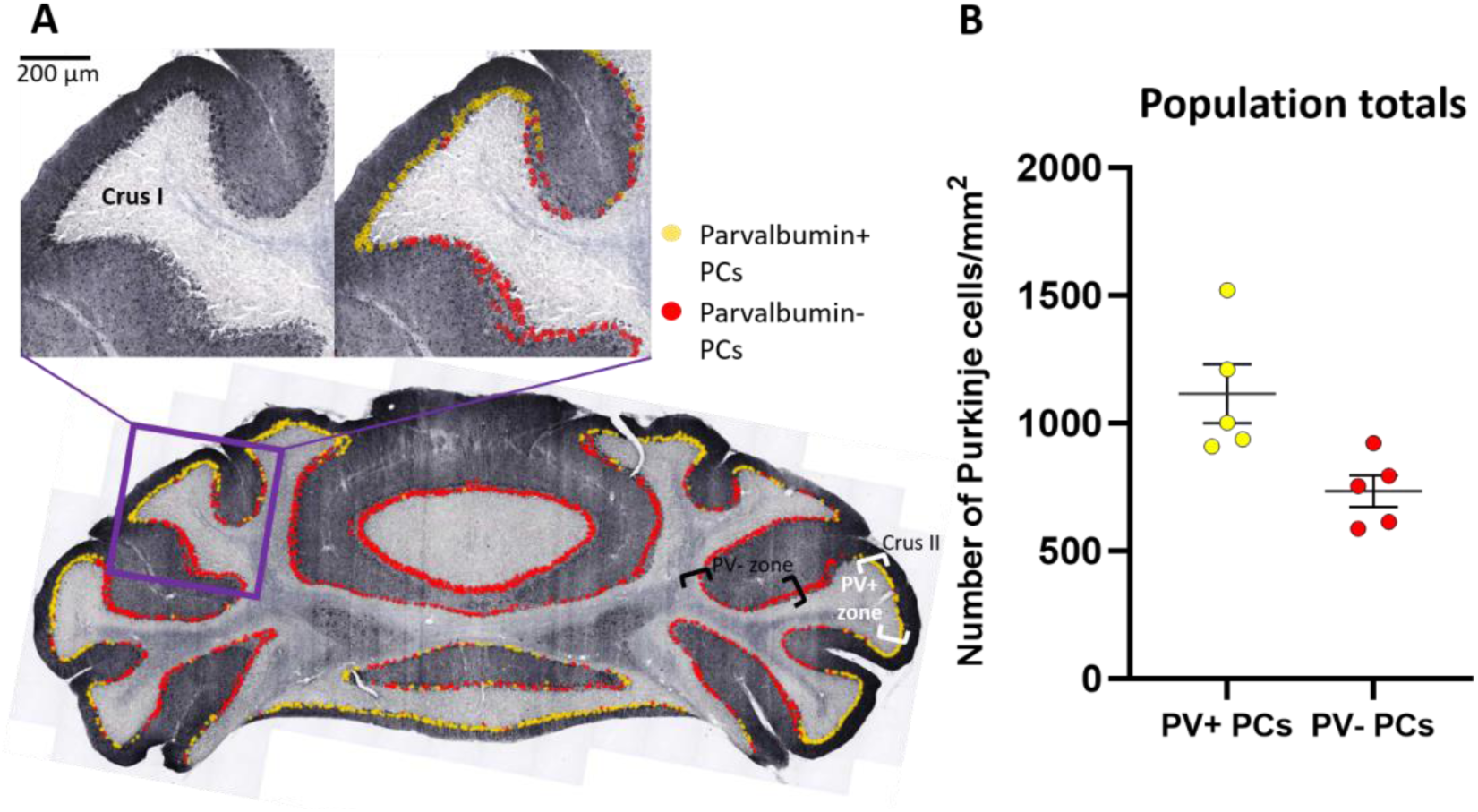
Parvalbumin-positive Purkinje cells segregate to the surface of the cerebellum within lobules. (A) An artificial intelligence model was built (Aiforia™) to analyze the distribution of parvalbumin-positive (yellow circles) and -negative (red circles) Purkinje cells throughout the cerebellum. Distribution of cells in serial sections throughout the entire cerebellum can be found in supplemental figure 1. (B) Purkinje cells positive for parvalbumin represent 60.3% of the total population while negative cells represent 39.7%. Total cells counted from each serial section through the cerebellum by the artificial intelligence model were divided by the area of the total Purkinje cell layer in five C57/Bl6 mice. Data are represented as mean ± SEM.

### Intrinsic firing properties of Purkinje cells are different across parvalbumin subtypes

As PCs displayed clear segregation of subtypes based on PV staining and PV is a well-known calcium buffer that can modify spike rates, we reasoned that PV could be an underlying mechanism for intrinsic firing differences that have been reported for aldolase C stripes. While Figure 3 shows the cerebellar vermis in order to clearly visualize aldolase C stripes and compare populations, PCs in cerebellar lobule crus II were chosen for electrophysiological recordings based on the clear segregation of PV from the AI distribution maps, although PV-cells could also be chosen from crus I at the border between crus 1 and crus II. Carefully choosing from the areas with yellow labeled (PV+) or red labeled (PV-) cells, as shown in Figure 4A, provided reasonable confidence as to the identity of each PC and keeping recording areas consistent in lobules crus I and crus II kept regional variation to a minimum. Unfortunately, after acquiring mice expressing tdTomato under the PV promoter, it was found that while most known PV+ cell types did express tdTomato throughout the brain (Kaiser et al., 2016), PCs did not have detectable tdTomato expression (supplemental figure 2). Consequently, this mouse strain is not useful for identifying PV+ PCs and could not be used to choose PCs for recordings. Additionally, due to the high expression of PV in dendrites and interneuron innervation to PC soma, post-staining the slices used for recording did not give reliable delineation of the cell types with fluorescence. For this reason, we recorded from a large number of cells to assess differences in firing properties between the two populations to minimize any impact of erroneous inclusion of aldolase C positive cells. It thus cannot be ruled out that aldolase C positive cells were recorded from in the generally PV positive areas, but both groups would have representation of aldolase C PCs.

Loose cell-attached recordings were obtained over a period of five minutes for each cell while blocking synaptic input so that only intrinsic firing properties were examined. PCs spontaneously display either tonic firing or burst-pause sequences (Llinas and Sugimori, 1980a,b; Chang et al., 1993; Womack and Khodakhah, 2002; Lowenstein et al., 2005; Oldfield et al., 2005; Yartsev et al., 2009; Wang et al., 2002; Williams et al., 2012; Cheron et al., 2014; Zhou et al., 2015). However, there is much diversity between PCs in terms of firing rate, time between bursts and length of bursts so that each cell displays a pattern that is quite different from its neighbors (Figure 5). The pattern that each cell displays is remarkably stable over time. A PC will maintain tonic firing at a relatively constant rate or will consistently burst and pause for at least several hours as reported in previous studies (Llinas and Sugimori 1980a; Hausser and Clark, 1997; Womack and Khodakhah, 2002) and confirmed in our laboratory (supplemental figure 3). This diversity in spike properties makes direct comparison more difficult so the spike rate was only assessed over the time the PC was actually firing within the five minutes total recording time (green bars in Figure 5). PCs in the PV+ areas typically spend less time in the bursting state (more time pausing) and are less likely to fire tonically (100% in figure) for the full five minutes, but when firing they tend to display a higher firing rate than PCs in the PV-areas (Figure 6).

**Figure 5.**
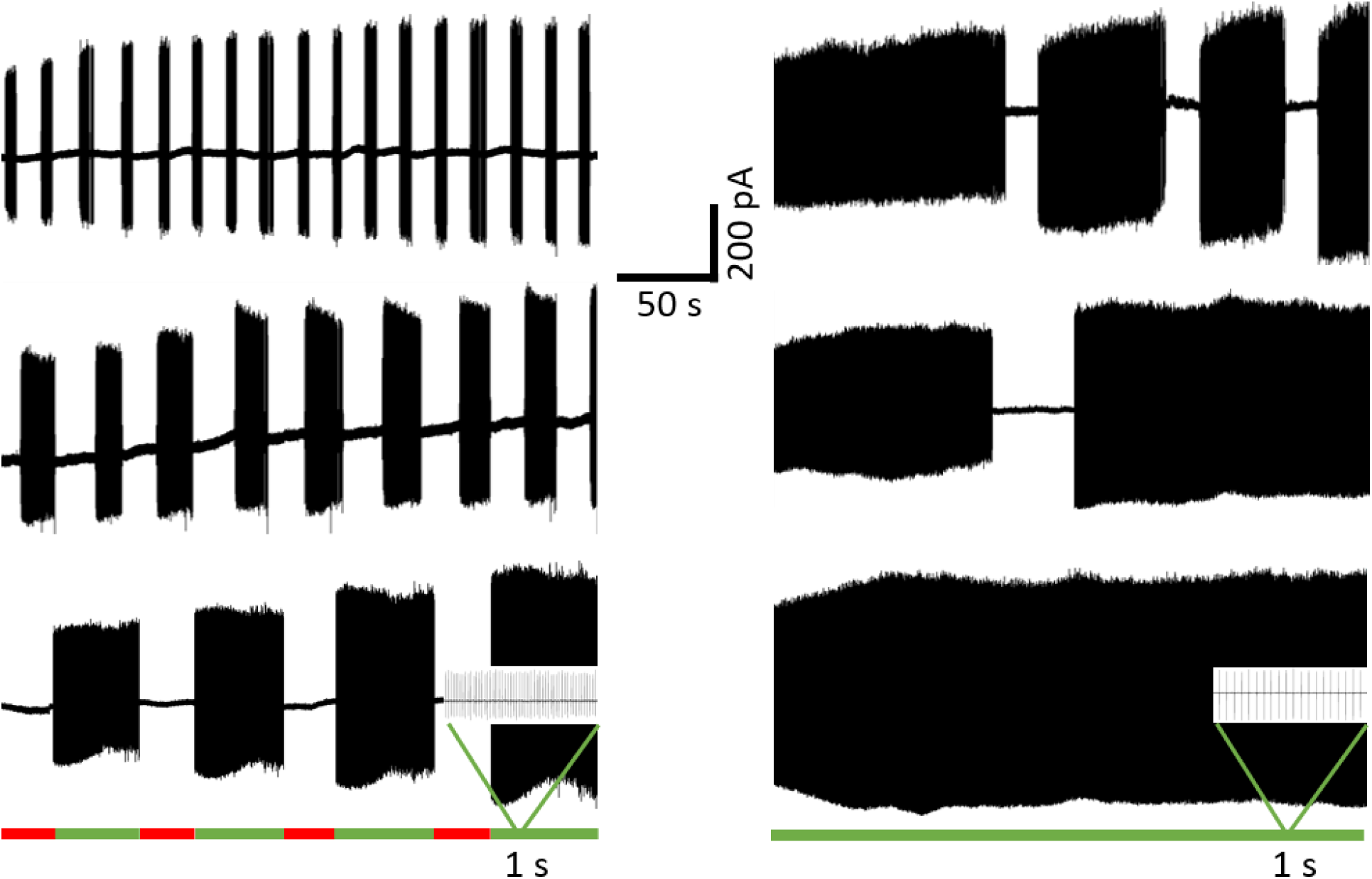
Diversity of Purkinje cell firing phenotypes. Cell-attached recordings over five minutes reveal the distinct firing patterns that individual neurons display, which continuously repeats over several hours. Burst firing displays various lengths of time of firing (green bars in lower panel example) and pausing (red bars). Individual spikes, as can be seen in the one second inset example of each lower trace, were counted with the software Neuromatic and divided by the total time spent firing (sum of green bars) for measures of firing rate in subsequent figures. Percentage of time firing is given as length of bursts divided by the total amount of time (always five minutes). An example of a tonic firing cell is in the lower right panel. Changes in amplitude are not relevant, as they are a well-known function of the strength of the loose seal (negative pressure applied) shifting over time.

**Figure 6.**
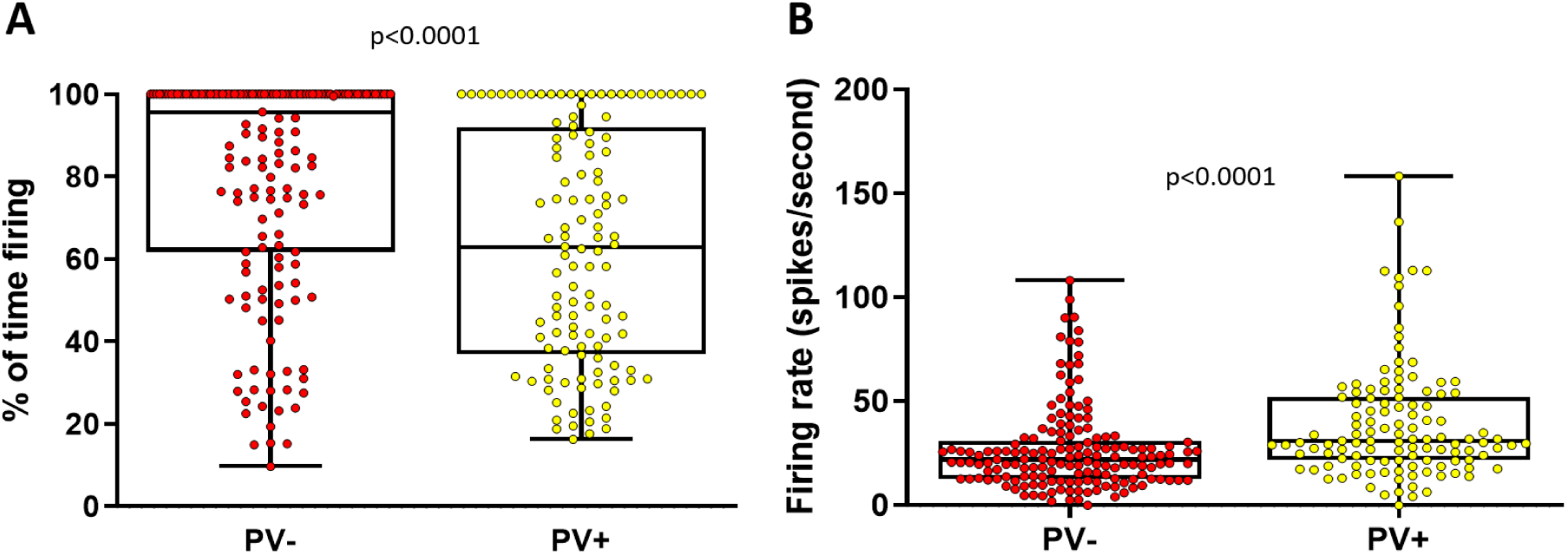
Areas containing parvalbumin-negative or -positive cells show differences in intrinsic firing properties. A) Over five minutes of cell-attached recording time, Purkinje cells within parvalbumin-negative zones spend more time in the “active” or firing state with more (n=80/164) cells tonically active (100% of time firing), while parvalbumin-positive cells had longer periods of pausing and less cells tonically active (n=25/121 cells). B) Additionally, while parvalbumin-positive cells tend to spend less time firing, during the firing bursts they typically display higher firing rates (n=10 mice). P-values were calculated with a Mann-Whitney test.

### Parvalbumin contributes to a higher spike rate and the burst-pause phenotype in Purkinje cell subtypes

To assess whether PV contributes to spike rate in PCs, cells from PV-areas were chosen and the baseline firing rate over five minutes was recorded before bath application of cell permeable EGTA/AM (150µM). EGTA is a calcium buffer with similar properties to PV (Caillard et al., 2000; Schwaller et al., 2002), although parvalbumin has additional properties at high concentrations (Eggermann and Jonas, 2013). We hypothesized that increased calcium buffering leads to an increased spike rate and mimicking PV buffering capacity with EGTA in PV-cells would lead to a detectable increase in firing rate. Indeed, addition of EGTA/AM resulted in a significant increase in firing rate (Figure 7A,B) in cells that otherwise fire at a very consistent rate over time (supplemental figure 3). Furthermore, pausing behavior increased in the EGTA/AM group (Figure 7C), which suggests sequestration of calcium by EGTA (and therefore PV) likely influences the ability of cells to maintain tonic firing.

**Figure 7.**
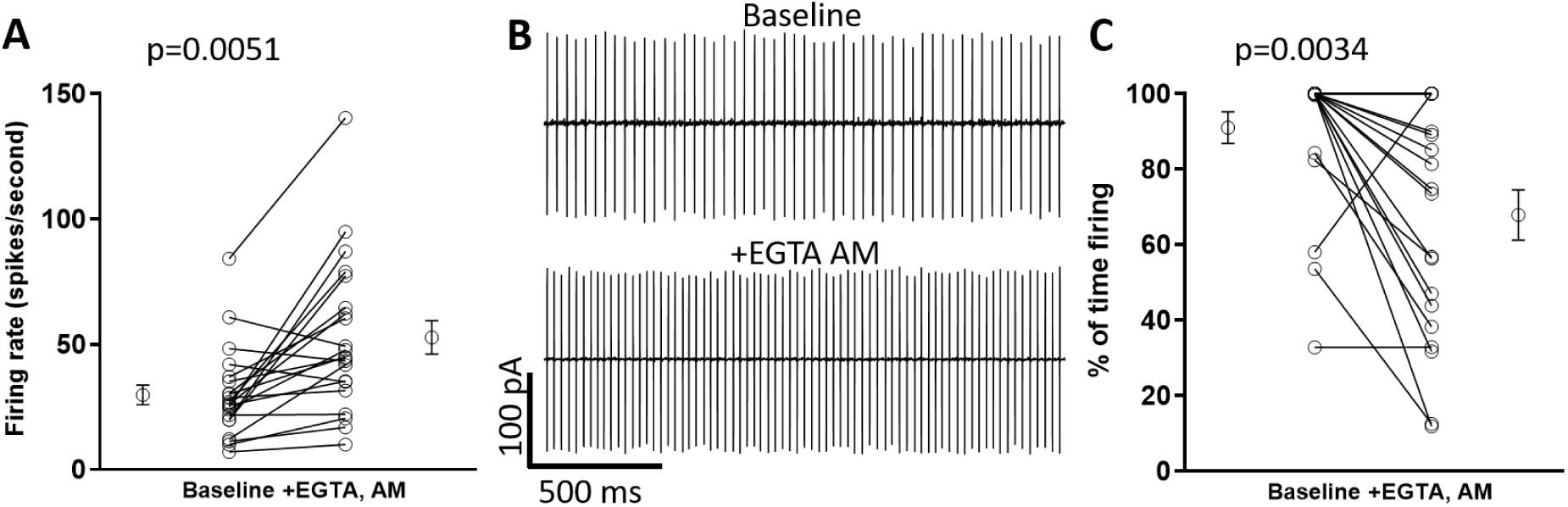
Cells in parvalbumin-negative regions increase firing rate and pausing time when exposed to a calcium buffer. (A) Purkinje cells were chosen from parvalbumin-negative regions and each firing rate was calculated over five minutes recording time. Bath application of cell permeable EGTA/AM, which shares properties of calcium buffering similar to parvalbumin, resulted in a significant increase in firing rate (paired t-test) within cells that would otherwise maintain a consistent firing rate over time (supplement figure 3). (B) Tonic firing rate over two seconds (to resolve individual spikes) increases after exposure to EGTA/AM. This suggests that parvalbumin is directly involved in regulating firing rate. C) The percentage of time spent firing decreased after application of EGTA/AM (Wilcoxon matched-pairs signed rank test). Therefore, tonically firing cells (100% active) began to display burst-pause behavior. Circles at 100% overlap in the graph; 5/15 tonically active cells remained tonically active while 10/15 cells began to pause. n=21 cells, 4 mice. The open circles with error bars on the sides in A and C indicate mean ±SEM.

PV’s contribution to spike rate was further confirmed by recording from PCs that no longer express PV. All cells in PV knock-out (KO) and littermate wild-type (WT) mice were chosen from the typically PV+ area of crus II, as in previous recordings. The PV KO PCs no longer displayed the full firing rate range or average mean expected for the PV+ zone, while the WT littermate recordings reproduced the range found in the previous experiment (Figure 8A). However, percentage of time firing did not significantly shift (Figure 8B). PV KO was confirmed with a western blot (supplemental figure 4).

**Figure 8.**
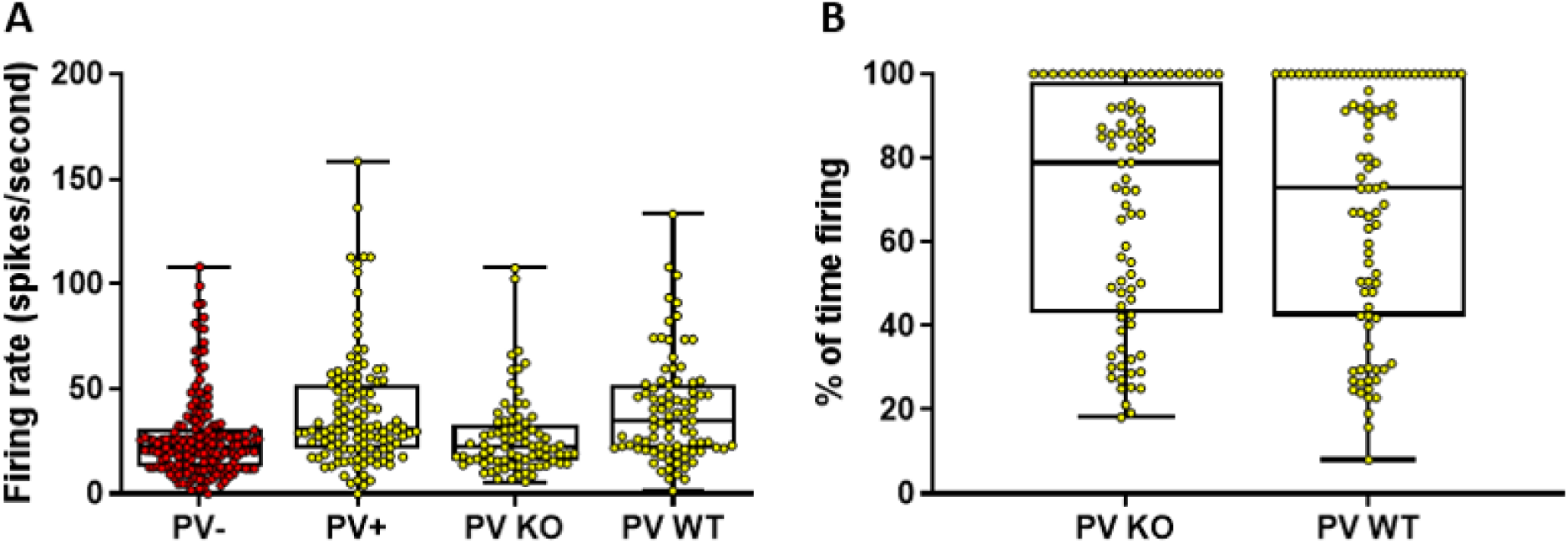
Parvalbumin knock-out mice have PCs with decreased firing rates. A) The PV- and PV+ columns for firing rate are reproduced from figure 5 to compare the PV KO mice. Over five minutes of cell-attached recording time, Purkinje cells in the typically PV+ zone of mice that no longer express PV have lower average firing rates that resemble PCs in PV-zones (n=80 PCs, 3 mice). Their wild-type littermates (PV WT) show the average firing rate as previously found for wild-type PV+ cells (n=86 PCs, 3 mice). B) PV KO mice do not show significant differences in percentage of time firing.

### Potassium and calcium channel blockade differentially affect Purkinje cell parvalbumin subtypes

As a group, cells exposed to EGTA/AM both increased firing rate and pausing time. However, individual cells did not always follow this pattern. For example, several cells did not increase firing rate and one cell increased active time, suggesting that other factors may also contribute to an individual cell’s bursting behavior and spike rate. Furthermore, the burst-pause behavior of PV KO cells did not shift as in the EGTA experiment, which likely reflects dependence on other factors than PV in this subpopulation. Burst-pause dynamics in PCs are known to depend on the interplay of sodium, calcium and potassium channels (Llinas and Sugimori, 1980; Raman and Bean, 1997, 1999; Nam and Hockberger, 1997; Womack and Khodakhah, 2002; Shim et al., 2018), which could also contribute to differences in intrinsic firing patterns between individual PCs. Therefore, we applied either 1mM TEA to block potassium channels or 500nM ω-Conotoxin MVIIC to block P/Q type calcium channels and both PV subtypes were assessed for differential responses (Figure 9).

**Figure 9.**
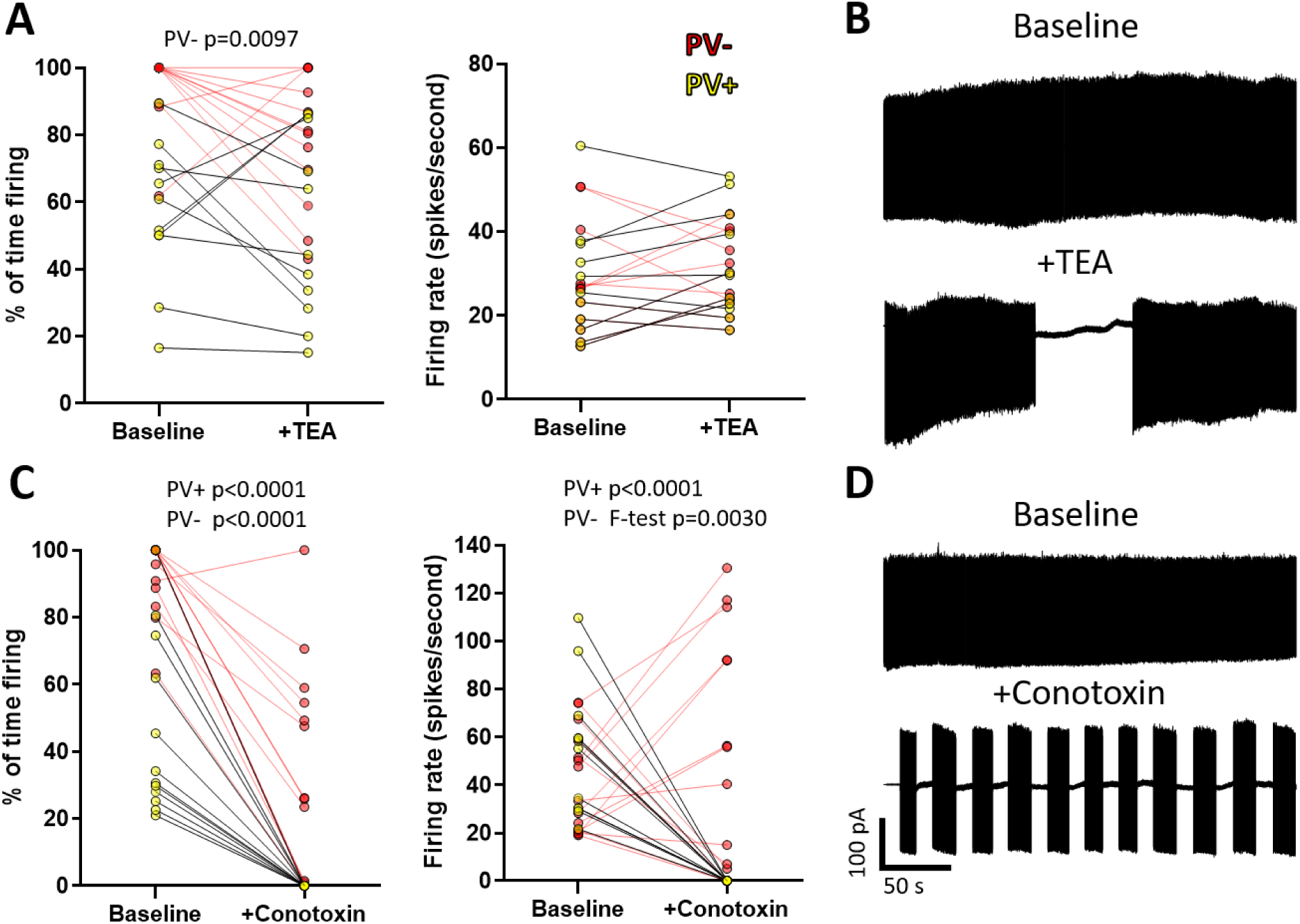
Cells in parvalbumin-positive and -negative regions show differential responses to potassium and calcium channel block. A) Application of 1mM TEA had variable effects on total time spent firing and firing rate over the five minute recording time, but consistently changed the firing pattern of parvalbumin-negative Purkinje cells through increased pausing time (Wilcoxon matched-pairs signed rank test; PV-n=12, PV+ n=11) B) For example, a tonically firing cell became a pausing cell. C) Application of 500nM ω-Conotoxin MVIIC reduced active time in both subtypes (Wilcoxon matched-pairs signed rank test), but caused all parvalbumin-positive cells (n=12) to go silent while several parvalbumin-negative cells (n=15) increased their firing rate during bursts, which increased the variance of the group (F-test). D) Similar to TEA, conotoxin produced distinct shifts in firing patterns that were variable from cell to cell.

Interestingly, both TEA and ω-Conotoxin MVICC application induced changes in the pattern of firing of PCs, but the particular response was associated with the PV subtype. PV+ PCs did not show significant differences in time spent firing or in firing rate, but PV-cells decreased time spent firing, which is reflective of the induced pausing behavior (Figure 9A). Qualitatively, blockade of potassium channels with TEA could completely transform the firing phenotype. For example a tonically firing cell could become a PC with burst-pause sequences (Figure 9B). Similarly, ω-Conotoxin MVICC application transformed tonically firing cells into bursting cells (Figure 9D). Each tonically firing cell could be shifted into a pausing cell, but the length of pauses and bursts produced were unpredictable, suggesting the overall firing phenotype is dictated by the combination of channel types. Blockade of P/Q type calcium channels consistently decreased the total time spent firing in PV-cells, but did not silence these cells. However, all PCs in PV+ areas were silenced by ω-Conotoxin MVICC application, driving their firing rate to zero and several PV-PCs displayed increased firing rates during periods of bursting, which increased the variance of the group firing rates.

Collectively, these different mechanisms that regulate and respond to intracellular calcium appear to contribute to the diverse spontaneous firing phenotypes displayed by PCs. PV expression can contribute to the higher firing rates and pausing behavior in individual PCs while both potassium channels and P/Q type calcium channels contribute to setting the burst-pause behavior, but exert variable effects on firing rate from cell to cell. Since PCs in the PV+ areas were all silenced with ω-Conotoxin application, it is likely this cell subtype is most reliant on P/Q type calcium channels to initiate burst-pause sequences, suggesting that PV may be highly expressed as a means to buffer increased calcium influx through the P/Q type channels. Cells in PV-areas also reduced time spent firing, but did not go silent. This suggests individual PCs regulate the composition of various channel types and intracellular calcium binding proteins to produce the diverse intrinsic firing phenotypes observed.

### Limitations

Due to the unexpected finding that PV-tdTomato did not express PV within Purkinje cells (it was present in interneurons) in any cerebellar area examined, confirmed PV+ cells could not be chosen for electrophysiological recordings and post-stain of PV with fluorescence was unreliable in the thick sections. Therefore, the identity of each cell could not be confirmed and both groups (PV+ and PV-) likely contain aldolase C positive PCs since striping occurs throughout crus II. Strategies to discriminate between the cell types in future studies may reduce variability between the two groups.

Recordings of PCs before and after application of TEA and ω-Conotoxin MVIIC revealed very distinct shifts in the overall firing phenotype observed, demonstrating their importance for the overall shape of the burst and pause times. However, whole-cell recordings will be useful to confirm differences in number and function of the various channel types for each cell and to tease apart the various contributions of potassium channel subtypes, as TEA will exert effects on all potassium channels.

PV knock-out mice and EGTA/AM were studied for their effect on firing rate, however, more specific genetic strategies for cell-specific knock-out and rescue of PV would be an important future study. Additionally, methods to quantify single-cell expression of PV in PCs will be useful since antibody titrations can affect the appearance of PCs and some may have very low expression instead of zero expression.

## Discussion

### Heterogeneous populations of Purkinje cells

The widely held assumption of an operationally uniform cerebellar cortex or “universal cerebellar transform” has come under recent scrutiny, as evidence increasingly points to variation in anatomy and physiology regionally (for review see: Cerminara et al., 2015). Many of the markers found to be differentially expressed across lobules label synaptic proteins, such as excitatory amino acid transporter (EAAT4) (Nagao et al., 1997; Dehnes et al., 1998), mGluRs (Mateos et al., 2012) and GABA receptors (Fritschy et al., 1999; Chung et al., 2008), among others. Differences in synaptic physiology would therefore be assumed to underlie the reported differences in firing properties between cerebellar modules. However, regional variation in simple spike frequency has been reported even when pharmacologically blocking all input (Zhou et al., 2014), indicating that intrinsic properties dictate PC firing phenotypes that are then modulated by synaptic input. The underlying mechanisms that drive these differences in PC spontaneous firing have so far been elusive. Our data provides evidence that individual PCs are diverse in their utilization of both ion channels and an intracellular calcium binding protein that can account, at least in part, for the differences observed in firing rate and burst-pause behavior that may dictate intrinsic excitability and thereby information coding to the deep cerebellar nuclei.

PV, long thought to be a general PC marker, is restricted to a subpopulation that does not display the typical striping phenotype. Instead, PV+ PCs are segregated to the surface (perimeter) of the cerebellum. This difference in PV expression contributes to the mounting evidence that PCs are divided into many subtypes based on expression of both intracellular and membrane proteins. For this reason, the common method of pooling all PCs when analyzing differences in a mutant animal, for example, may obscure important differences due to the substantial heterogeneity across these cells. More precisely defining which groups of PCs to examine may therefore provide better insight into potential differences.

### Intrinsic differences of parvalbumin subpopulations

The segregation of PV positive PCs to the surface of the cerebellum correlates with the location of cells that often display the burst-pause phenotype instead of tonic firing. This observation has been reported in the literature, where Zhou et al. (2015) methodically recorded from PCs across the cerebellum and also found differences in burst-pause firing based on location, stating that PCs with long pauses segregate to the surface of the cerebellum while non-pausing cells tend to have an interior location. As our PV maps precisely correspond to these phenotypes, PV+ PCs appear to be a distinct subpopulation. Differences in expression of other molecules within the PV populations is likely, which future studies may illuminate.

The present data suggests that ion channel expression is variable across PC PV subtypes, as the contribution of various ion channels assessed by pharmacological blockade is different depending on which PC subtype is examined. It is well documented that burst-pause dynamics are controlled by the interaction of voltage gated sodium, calcium and potassium channels (Llinas and Sugimori, 1980; Raman and Bean, 1997, 1999; Nam and Hockberger, 1997; Womack and Khodakhah, 2002; Shim et al., 2018), but the present data suggests that these interactions can vary from cell to cell. Blocking P/Q type calcium channels resulted in silencing of PV+ PCs over the time course examined, while block of PV-PCs resulted in longer pause sequences, but often increased firing rate. Strikingly, calcium channel block also shifted tonically firing cells into the burst-pause phenotype. An informative study (Womack and Khodakhah, 2002) examining burst-pause dynamics in PCs also used ω-Conotoxin MVIIC over a similar time course to block P/Q type calcium channels and observed that bursting cells go silent after application, while tonically active cells increased their firing rate, burst and then went silent. We also observed this behavior, but it was restricted to the PV+ regions. Although there are not as many tonically active cells in this area, they are present. The tonically active cells in the PV+ region went silent with ω-Conotoxin MVIIC application, while the tonically active cells in the PV-region were not silenced over the course of forty-five minutes. As the Womack and Khodakhah, 2002 study was performed in the cerebellar vermis, direct comparisons are difficult and future work is needed to show whether PV+ and PV-regions in the vermis show the same phenotype as those in the cerebellar hemisphere.

Potassium channel blockade only slightly increased or decreased firing rate in individual cells and unpredictably altered the burst-pause dynamics observed, suggesting that individual cells may regulate the number of channels expressed and/or differentially modify potassium channel function to contribute to the distinct firing patterns observed. McKay and Turner (2004) delineated the role of potassium channels in the bursting behavior of PCs in the rat vermis and found that application of TEA could shift the pattern of bursting and bursting frequency, which they reported to mainly be dependent on Kv3 channels. We observed a similar shift in PV-cells (reflected in time spent firing), but the PV+ cell group mean did not significantly shift, suggesting that these differences may be subtype-dependent, but may also be due to regional differences. Interestingly, Kv3 channels have been reported to have high variability of expression in PCs from cell to cell (Veys et al., 2013), which could explain the various responses we observed. This variability may be a promising avenue for exploration into more precise characterization of individual PC phenotypes.

Potassium channels in PCs are calcium sensitive and regulate both sodium and calcium spike discharge (McKay and Turner, 2004) while P/Q type calcium channels pass calcium directly (Llinas and Sugimori, 1980), therefore, calcium utilization and regulation is crucial to the generation of bursting behavior in PCs. Therefore, it is reasonable to suspect differences in ion channel function and/or expression as a mechanism underlying differences in firing rate and bursting properties between individual cells. Block of both potassium and calcium channels caused the PC to shift into different patterns of firing. This is interesting because the pattern is usually very stable over time and could be changed abruptly to a new firing phenotype after blockers were added. Furthermore, the bursting did not become random, it shifted into a new stable pattern after the addition of blockers, but the pattern was unpredictable in each cell. This suggests ion channels are not expressed equally or functioning in the same way between neighboring cells.

In addition to these ionic mechanisms that can control intrinsic excitability, calcium buffers likely add further complexity by modifying the amount of calcium available and, consequentially, confer the capacity for higher spike rates and sustained firing. We show that mimic of PV calcium buffering with EGTA in PV-areas results in increased firing rate and increased pausing behavior that resembles the cells in PV+ areas, suggesting a direct role of PV in the intrinsic firing differences observed between subpopulations.

### Implications for intrinsic plasticity

As all synaptic contribution was blocked in our preparation, the phenotypes recorded represent spontaneous/intrinsic properties of individual PCs and are highly diverse from cell to cell based on length of burst as well as firing rates during bursts or tonic activity. It is unclear to what extent these spontaneous properties can shift within one cell over time. During development, PCs have been reported to fire more irregularly than they do after four weeks of maturity (Arancillo et al., 2015), suggesting that phenotypes are acquired due to development of connectivity and activity-dependent processes. An emerging concept particularly relevant for PCs is that information storage may not only depend on synaptic plasticity, but on activity-dependent modulation of intrinsic excitability, or intrinsic plasticity, as ion channels are modified based on input (for review see: Shim et al., 2018). PC membrane bistability, characterized by burst-pause sequences, can be modified by climbing fiber input (Cheron et al., 2014), hyperpolarization-activated mixed cationic current IH (Williams et al., 2002), Bergmann glia (Wang et al., 2012) and interneurons (Oldfield et al., 2010). As our data provide evidence that alterations in PV expression as well as potassium and calcium channel function can shift cells between different firing phenotypes, it is likely these mechanisms are utilized by PCs for intrinsic plasticity and may be important for the incorporation of functionally distinct PCs in behavior, as recently shown in zebrafish locomotor tasks (Chang et al., 2020).

### Summary

In sum, we provide evidence for calcium homeostasis as the underlying driver of the diversity in PC firing phenotypes and demonstrate a novel route for labeling PC subpopulations. Parvalbumin is able to affect intrinsic firing rate and bursting behavior, but differences in ion channel expression are likely the major drivers of diversity in the shape of the burst-pause behavior of PCs. Parvalbumin, as a marker, provides researchers new tools for fine-mapping of cerebellar microcircuitry and emphasizes the importance of single-cell studies, as PCs are diverse in their molecular profile and firing phenotypes. Since baseline differences in spontaneous firing of PCs provide coding information to the deep cerebellar nuclei that can be modified by synaptic input arriving to each PC, future research may help to clarify the relationship between intrinsic PC coding and synaptic modification that results in signaling to output structures. This may serve to clarify how these differences in PC subtype may enable information coding across cerebellar microzones and modules, as well as their impact for the many conditions represented by cerebellar processing deficits.

## Acknowledgements

The authors acknowledge helpful assistance from Dr. Madel Durens and Dr. Gloria Hoffman. We thank Dr. John P. Hussman for his critical review of the manuscript. This work is supported by Hussman Foundation grants HIAS #15001 and HIAS #17001 to GJB and Hussman Foundation grant HIAS #18001 to SH.

## Author contributions

Conceptualization, C.B.; Methodology, C.B., L.A.S, M.K., M.B. and S.H.; Investigation, C.B.; Writing – Original Draft, C.B.; Writing – Review & Editing, C.B., L.A.S., M.K., M.S.B., S.H. and G.J.B.; Funding Acquisition, S.H. and G.J.B.; Resources, C.B., L.A.S., Y-C.L., S.H. and G.J.B.; Supervision, S.H. and G.J.B

---

**Supplemental Figure 1.**
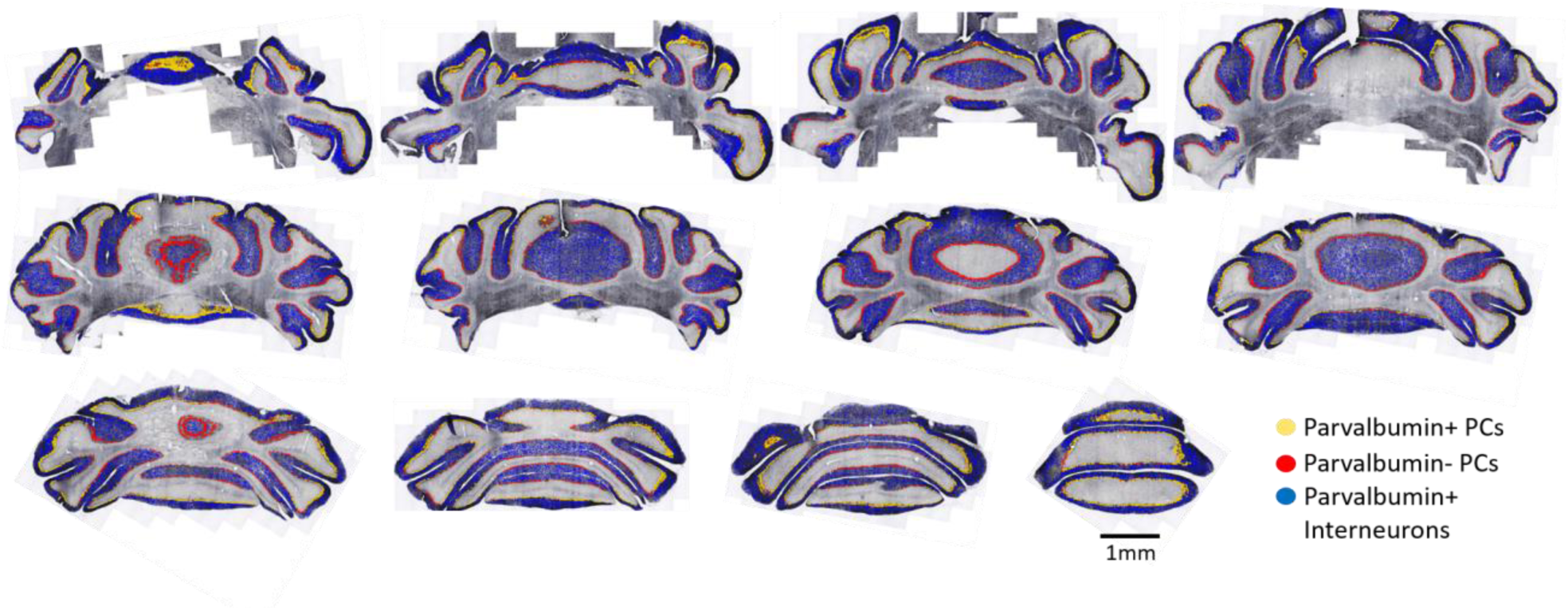
Distribution of parvalbumin-positive cells throughout the cerebellum. Serial (every 6^th^ section) 40µm coronal sections through C57/Bl6 cerebellum stained for parvalbumin. Images are shown left to right in rostral to caudal order. Yellow circles represent parvalbumin-positive Purkinje cells, red represent parvalbumin-negative Purkinje cells and blue circles represent interneurons recognized by a customized artificial intelligence algorithm (Aiforia™). Positive Purkinje cells continue to segregate toward the surface of the cerebellum in each plane.

**Supplemental Figure 2.**
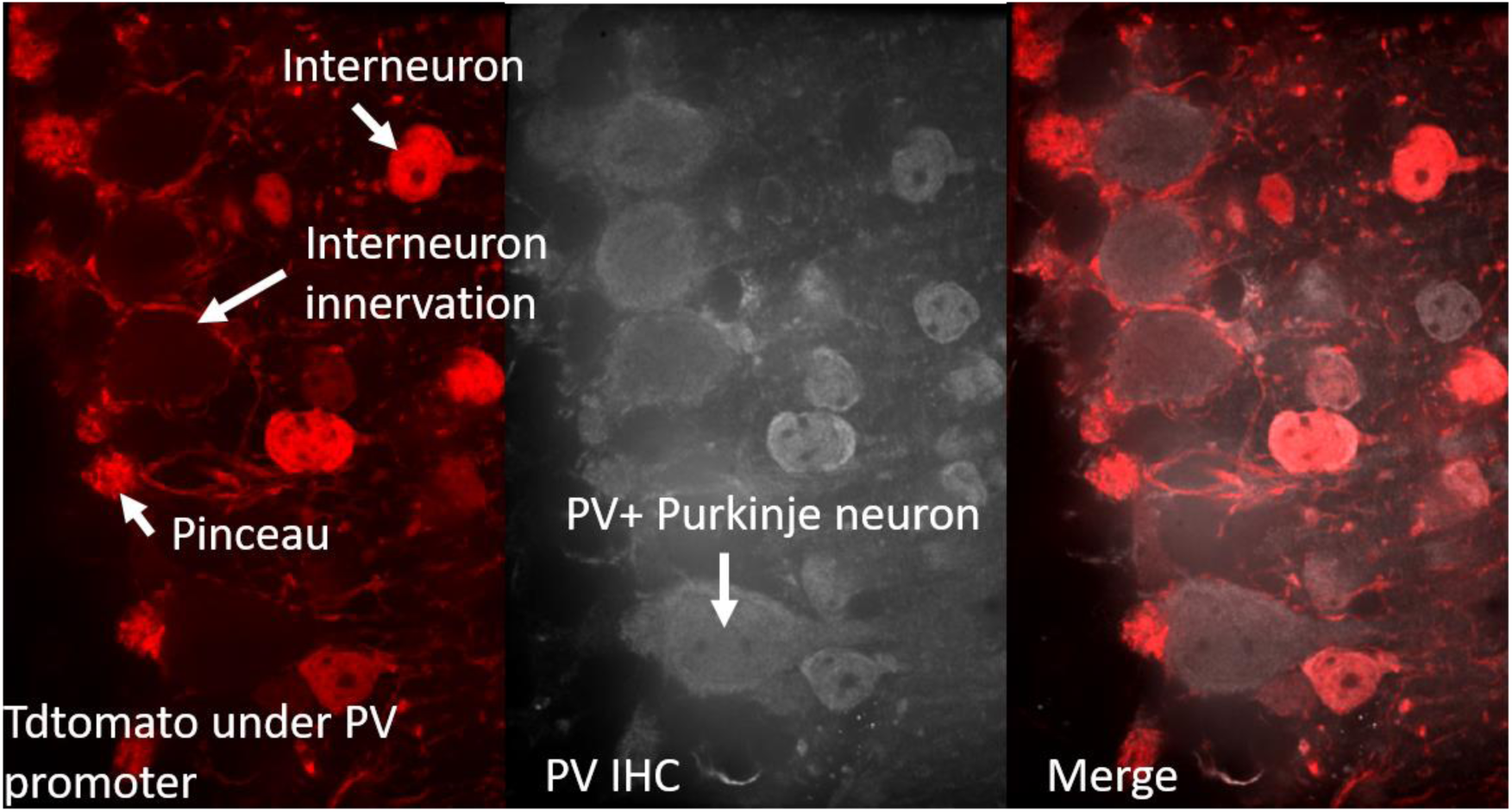
The PV-tdTomato mouse line does not label Purkinje cells. TdTomato can be observed within interneurons in the molecular layer, which innervate the Purkinje cell soma forming an outer ring along with the pinceau, but expression is not observed intracellularly in Purkinje cells in any region examined. Parvalbumin immunohistochemistry, however, does label intracellular parvalbumin within Purkinje cells. Therefore, this mouse line does not allow for targeting parvalbumin positive Purkinje cells for electrophysiological recordings.

**Supplemental Figure 3.**
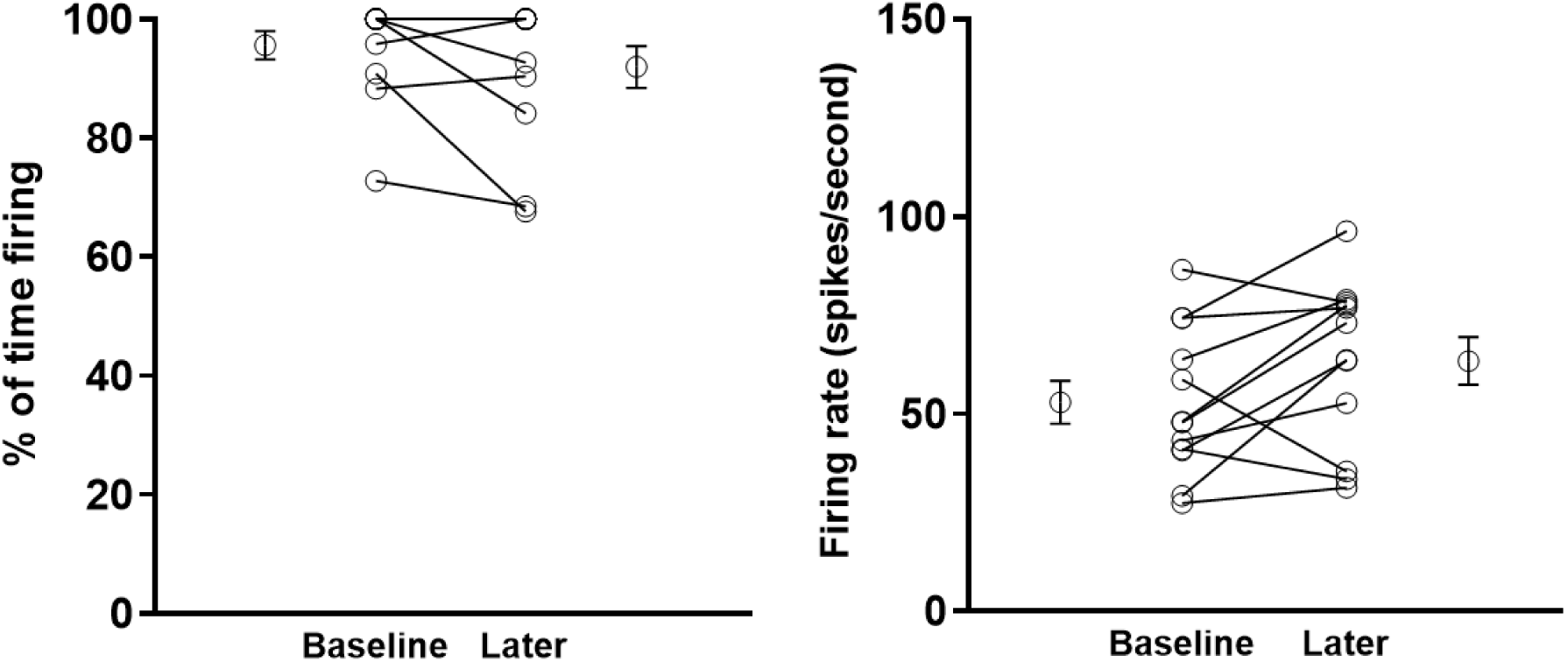
Purkinje cells maintain consistent firing patterns and rates over several hours. Recordings of individual Purkinje cells were performed over 1-3 hours without significant variation in percentage of time firing or firing rate of the last recording compared to the first. Both percentage of time firing and firing rate were recorded over the last five minutes of recording time and compared to the first five minutes. n=12 cells, 6/12 cells overlap at 100% to 100% in the left panel. No significant differences with a Wilcoxon matched-pairs signed rank test. The open circles with error bars on sides indicate mean ± SEM.

**Supplemental Figure 4.**
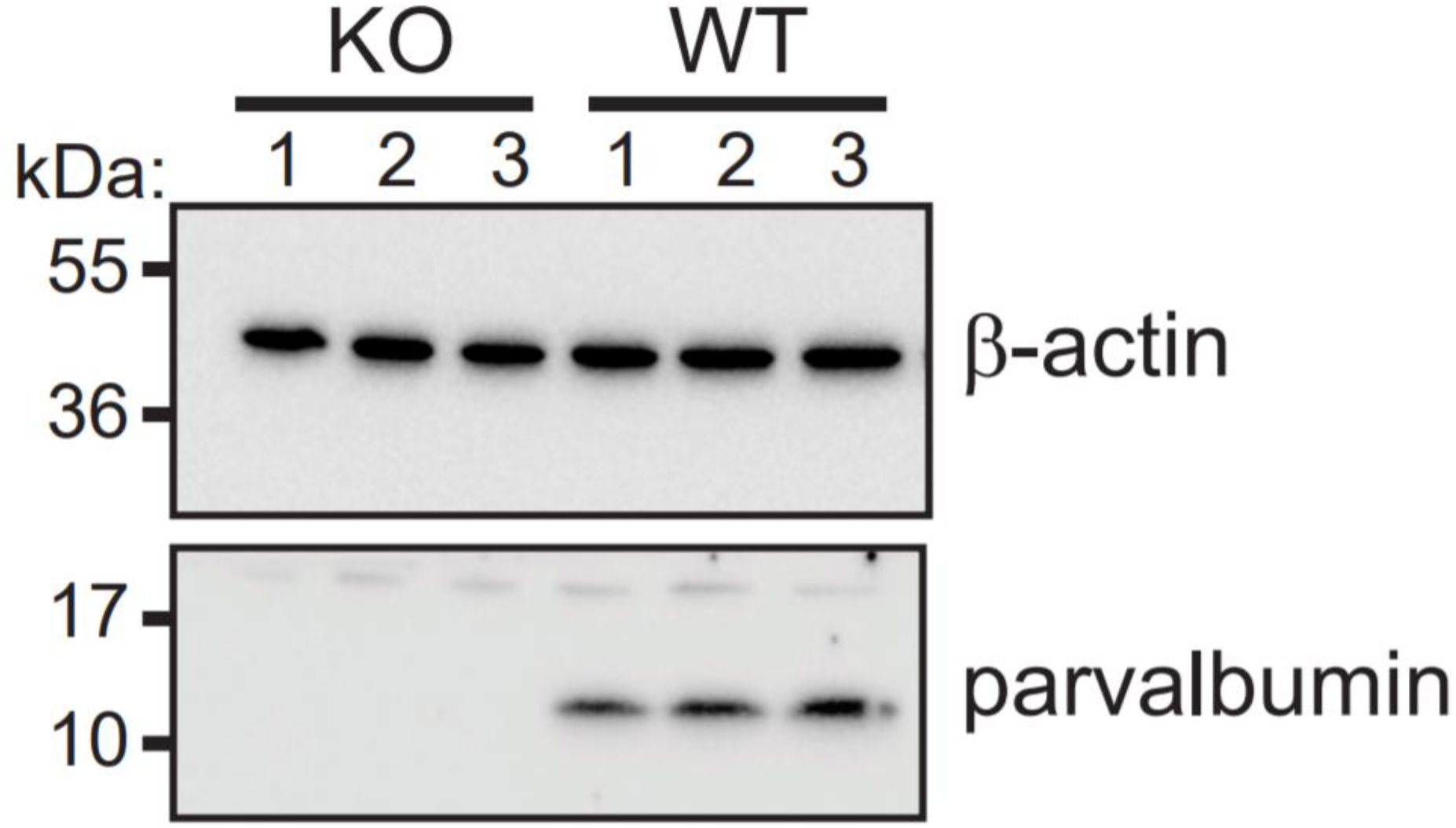
Confirmation of absence of parvalbumin expression in parvalbumin knock-out animals. Cropped image of immunoblot showing parvalbumin expression in all wild-type littermates but not in PV KO animals. β-actin was used as loading control. Membrane was cut at the 28kDa marker and pieces were blotted separately with indicated antibodies. N = 3 animals/genotype.

